# Quantitative metagenomics reveals fine-scale population dynamics across bacteria, archaea, and microbial eukaryotes in an estuarine–coastal continuum

**DOI:** 10.64898/2026.05.28.728481

**Authors:** Liang Zhao, Nathaniel P. Curtis, Ryan W. Paerl, Scott M. Gifford

## Abstract

Microbial communities are foundational to marine ecosystem function, yet their diversity is often obscured by broad taxonomic groupings and relative-abundance surveys that mask the dynamics of individual populations. This limitation is especially important across estuarine–coastal gradients, where microbial standing stocks, environmental conditions, and community composition vary sharply over space and time. Here, we used quantitative, genome-resolved metagenomics to examine microbial population dynamics across a one-year estuary-to-ocean transect spanning the Neuse River Estuary, Pamlico Sound, and adjacent North Atlantic shelf waters. Internal standard normalization enabled absolute abundance estimates for single-copy genes and metagenome-assembled genomes (MAGs), allowing individual populations to be tracked as genome equivalents per liter. Bacterial standing stocks were higher in estuarine waters, and communities varied primarily with salinity and season. We recovered 415 MAGs, including 386 bacterial genomes that represented, on average, 52% of bacterial genome equivalents, along with archaeal and eukaryotic representatives. Many abundant MAGs lacked close reference genomes, demonstrating that numerically important coastal populations remain poorly characterized. Genome-resolved abundances revealed pronounced niche partitioning among closely related taxa, including seasonal and spatial turnover of *Synechococcus*, *Cyanobium*, and *Vulcanococcus* populations associated with distinct pigment-defined cytometric groups. Rhodobacteraceae MAGs also showed population-specific correlations with picoeukaryotic MAGs, including a winter offshore *Planktomarina* population that reached 18% of total bacterial genome equivalents during a *Micromonas*-associated bloom. By providing absolute population abundances, this study transformed coastal microbiome surveys into numerical frameworks for resolving microbial population structure, ecological interactions, and biogeochemical relevance across dynamic environmental gradients.

**IMPORTANCE:** Estuarine and coastal waters contain diverse microbial communities that help regulate food webs and the cycling of carbon and nutrients, but many of the individual microbial populations responsible for these processes remain poorly understood. In this study, we examined bacteria, archaea, and small algae across the Neuse River Estuary, Pamlico Sound, and nearby coastal ocean waters over one year. By measuring the absolute abundance of individual microbial genomes, rather than only their relative proportions, we showed that closely related populations can have very different seasonal and spatial patterns. This was especially clear for cyanobacteria related to Synechococcus and heterotrophic bacteria in the family Rhodobacteraceae, which showed distinct population dynamics and associations with small algae. These results demonstrate how quantitative genome-resolved measurements can reveal hidden microbial population structure and improve our understanding of how microorganisms shape coastal ecosystem function.

## INTRODUCTION

Microorganisms form the foundation of marine ecosystems, driving biogeochemical cycles and sustaining higher trophic levels through primary production and recycling of dissolved organic matter (Azam and Malfatti, 2007; Moran, 2015). Despite their ecological importance, many microbial communities remain “black boxes” of which individual populations are not well characterized, including their metabolic potential and spatial and temporal dynamics. Understanding microbial composition and functional potential is especially crucial in productive coastal regions susceptible to anthropogenic perturbations such as eutrophication and habitat alterations, and where large human populations reside (Vitousek et al., 1997; Paerl et al., 2010).

Additionally, genomic characterizations of microbial communities usually provide only relative abundances, making cross-sample comparisons challenging because changes in relative abundance do not necessarily reflect changes in cell abundance. This limitation is especially important when comparing estuarine and coastal ecosystems, where microbial cell densities can vary by orders of magnitude and relative abundance alone can obscure whether individual populations are increasing, decreasing, or simply changing proportionally. The application of internal standard normalization techniques enables read counts to be converted into volumetric estimates of genome or gene concentrations in environmental samples, thereby moving beyond relative community composition to quantify the absolute standing stocks of individual microbial populations (Gifford et al., 2011, 2020; Moran et al., 2013; Satinsky et al., 2013; Lin et al., 2019; Sharpe et al., 2023, Bei et al., 2025).

The Albemarle-Pamlico Estuarine System (APES) is the largest lagoonal estuary in the United States and is characterized by shallow depths, low tidal exchange, and long water residence times, making it highly sensitive to freshwater inputs and seasonal hydrological variability (Paerl et al., 2009, 2018). These hydrological dynamics strongly influence nutrient availability, phytoplankton biomass, and phytoplankton community composition across the system (Pinckney et al., 1998; Gong et al., 2018, 2020; Sanchez-Gallego et al., 2025). In contrast, nearby North Atlantic continental shelf waters are generally less productive than APES waters and support distinct microbial communities (Wang et al., 2019). Long-term coastal monitoring has further shown that microbial communities in this region undergo pronounced seasonal shifts, including turnover among closely related taxa (Ward et al., 2017). Together, these features make the APES-to-coastal ocean continuum a useful natural gradient for examining how microbial populations vary across strong spatial and seasonal environmental gradients.

The hydrology and phytoplankton ecology of APES and adjacent North Atlantic shelf waters have been extensively studied through long-term monitoring, pigment analyses, microscopy, flow cytometry, and amplicon sequencing (Paerl et al., 2009, 2010; Peierls and Paerl, 2010; Ward et al., 2017; Gong et al., 2018, 2020; Wang et al., 2019; Johnson et al., 2025). However, comparatively little is known about the genome-resolved diversity, abundance, and dynamics of bacterioplankton populations in this system. This gap is especially important for ecologically significant groups such as *Synechococcus*, which are key primary producers in estuarine and coastal waters and exhibit seasonal changes in abundance, composition, and pigment phenotype, yet remain poorly resolved at the population-genomic level in APES and nearby coastal waters (Ward et al., 2017; Wang et al., 2019; Paerl et al., 2020; Johnson et al., 2025; Sanchez-Gallego et al., 2025). Similarly, the population-level dynamics of abundant heterotrophic bacterial taxa remain under-characterized, limiting our ability to connect microbial community structure with ecosystem processes across the estuarine-to-coastal gradient.

To improve our understanding of marine microbial population dynamics and their influence on coastal ecosystems, we conducted a quantitative metagenomics survey across the estuary-to-ocean continuum of the APES. We specifically examined: (1) how microbial communities changed with time and space, (2) the environmental drivers of composition changes, and (3) the distributions of individual genome-delineated bacterial populations of ecological significance. Surface metagenome samples were collected from three sampling sites representing Neuse River Estuary, Pamlico Sound and North Atlantic continental shelf (Onslow Bay) waters over a one-year period. Internal standard normalization was applied to estimate the volumetric abundances of single-copy genes and metagenome-assembled genomes (MAGs). Our results indicate that both community composition and the distributions of individual genome-delineated populations show strong spatial and seasonal patterns across the estuarine-to-coastal gradient. By coupling MAG reconstruction with internal standard normalization, we quantitatively resolved microbial population structure at genome-level resolution, revealing patterns that would be obscured by relative-abundance or higher-level taxonomic analyses alone.

## METHODS

### Field sample collection

Samples were collected from three sites: Pamlico Sound at 35.1225° N, −76.2006° W (PS5), the Neuse River Estuary at 35.0641° N, −76.5260° W (NRE180), and Onslow Bay at 34.6094° N, −76.6771° W (OB1) from Aug 2021 to Aug 2022 (Fig. 1A and 1B). Sample collection at PS5 and NRE180 was conducted in conjunction with the ModMon and FerryMon projects (Luettich et al., 2000; Buzzelli et al., 2002; Paerl et al., 2009, 2010). Surface seawater at approximately 1 m depth was prefiltered with a 250 µm mesh and collected into 2L bottles at the sampling site via diaphragm pump. The bottles were transported back to shore and filtered, with about 30 mins between water collection and the start of filtering. Approximately 2L of water was passed through a 3 µm pore polycarbonate pre-filter (142 mm diameter) and a 0.2 µm pore PES filter (142 mm diameter) to collect cells, and then the filters were flash frozen in liquid nitrogen. The volume of seawater filtered was recorded. Average filtering time was 15 minutes.

**Figure 1.**
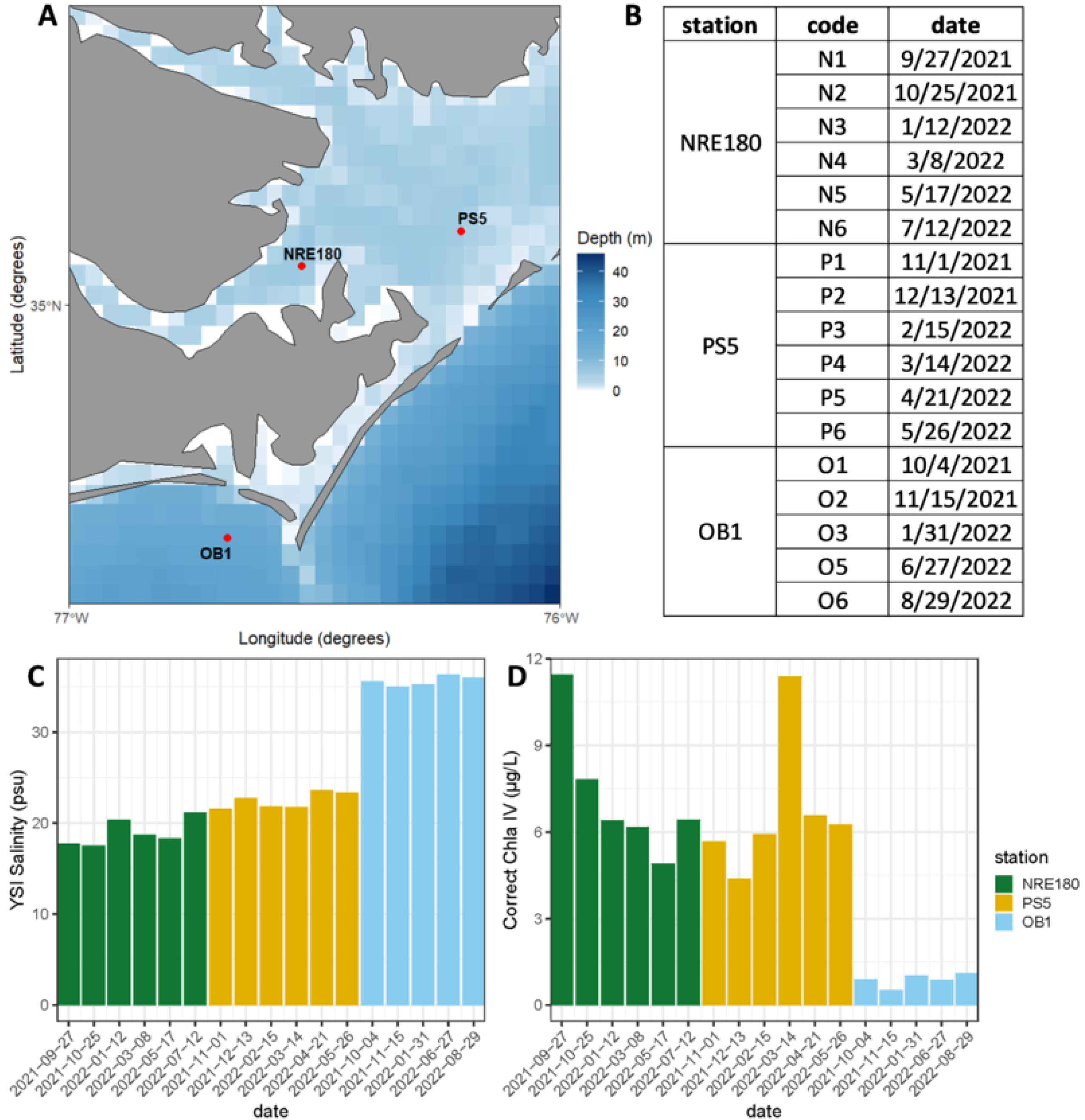
Study area, sampling timeline, and environmental gradients in the Albemarle-Pamlico Estuarine System (APES). (A) Map of sampling stations indicating the three primary sites along the estuary-to-ocean continuum: the lower Neuse River Estuary (NRE180), Pamlico Sound (PS5), and Onslow Bay (OB1). Bathymetry is represented by the blue color gradient indicating depth in meters. (B) Tabular summary of sampling trips, identifying the station codes and specific dates for the 17 quantitative metagenomic events spanning an annual cycle from September 2021 to August 2022. (C) Surface water salinity (psu) and D) Chlorophyll-a (µg/L) concentrations at each sampling point.

### DNA extraction and sequencing

Seventeen of the 0.2 µm filters were extracted for DNA with the DNeasy PowerMax Soil Kit (Qiagen) following the kit protocol. Three DNA internal standards (genome molecules of *Thermus thermophilus, Blautia producta*, and *Deinococcus radiodurans*) were each added at 2 ng each in 15 μL volumes to the bead tubes right before extraction (Gifford et al., 2020). Sequencing libraries and 250 bp Paired-End sequencing data were generated using the Illumina NovaSeq platform (San Diego, California, USA) at UNC’s High-Throughput Sequencing Facility (Chapel Hill, North Carolina, USA).

Trimmomatic was used to remove adaptor sequences and low-quality base pairs from both the forward and reverse reads with a sliding window of 10 bp and quality score cutoff of 20 (Bolger et al., 2014). Trimmed reads <50 bp were removed. Paired forward and reverse reads after quality control were merged using PEAR (default parameters) (Zhang et al., 2014). Internal standard reads were identified via a BLASTn search against the three internal standard genomes (BLAST passing criteria: *e*-value < 0.001, percent ID > 95%, alignment length > 50% of the read length, bit score > 50; genome IMG accession numbers: *T. thermophilus* [637000322], *D. radiodurans* [2556921628], *B. producta* [2515154176]). The identified internal standard reads were further annotated via a BLASTx (*e*-value < 0.001) homology search against the internal standard protein sequences, with bit score cutoff of 40 and percent identity cutoff of 95%. The reads identified as originating from the internal standard genomes were used to calculate recovery ratios and were removed from the downstream analyses. After trimming, merging, and removal of internal standard reads we retained a mean of 41 million reads per sample (range 32–57 million) (SI Table 2).

### Volumetric abundance estimation

To estimate gene and genome volumetric abundances, internal standard (IS) recovery ratios were calculated for both genes and genomes using methods described previously (Gifford et al., 2020; Sharpe et al., 2023). The relationship between the number of IS genomes added to the sample and the number of IS reads recovered were used to estimate volumetric abundances of genes in the sequence library. In this study, we estimated volumetric genome equivalent abundances using the single copy gene *recA*. The relationship between the number of IS genomes added to the sample and the number of IS genomes recovered was used to estimate volumetric abundances of MAGs based on the number of MAGs sequenced.

To estimate archaeal absolute abundance, we established a station-specific relationship between SingleM coverage and genome equivalents using bacterial data. For each station, we performed linear regression between bacterial SingleM coverage values and bacterial *recA* gene counts, with the y-intercept constrained to zero. The slope of each regression was used as a scaling factor to convert SingleM archaeal coverage values into estimates of archaeal genome equivalents.

### Taxonomic annotations

Reads were annotated with NCBI RefSeq (v.223; bacterial, eukaryotic, archaeal, and viral reference sequences) using DIAMOND blastx (Buchfink et al., 2015). A RecA protein database was assembled containing proteins from the RefSeq protein database annotated with the key words “recombinase RecA,” “protein RecA,” “recombinase A,” or “RecA protein.” Bacterial *recA* reads in the metagenomes were first identified by a DIAMOND homology search against this custom RecA protein database. Then the *recA* reads were re-annotated with RecA reference sequences from GTDB representative genomes (R202; TIGR2012) to match RecA taxonomy with MAG taxonomy (Parks et al., 2022). For archaeal annotations, the SingleM “pipe” workflow was used to annotate all samples and the archaeal annotations were extracted to visualize the archaeal compositions (Woodcroft et al., 2024). For eukaryotic annotations, reads were annotated with PhyloDB (https://github.com/allenlab/PhyloDB) and NCBI Refseq using DIAMOND blastx.

### Metagenome assembly and metagenome-assembled genomes (MAG) recovery

Paired and unpaired forward and reverse reads were assembled with SPAdes for individual samples (version 3.13; --meta and --only-assembler settings with default k-mer sizes: 21, 33, 55, 77) (Nurk et al., 2017). For binning, reads were mapped to contigs with Bowtie2 (version 2.3.4.1; default parameters) and the alignments were sorted and indexed using Samtools (version 1.9; default parameters). Contigs were binned with MetaBAT (version 2.13; contig size cutoff 2500bp), MaxBin2 (version 2.2.6; contig size cutoff 1000bp) and CONCOCT (version 1.1.0; contig size cutoff 1000bp) independently and the resulting bins were refined by DAS Tool (version 1.1.1; default parameters). dRep was used to dereplicate dastool bins resulting from all 17 assemblies with a 95% ANI threshold (Olm et al., 2017). The quality of the MAGs was assessed using CheckM (version 1.0.13) and taxonomy was assigned by GTDB-Tk (version 0.3.2; Genome Taxonomy Database release 04-RS89) (Chaumeil et al., 2020). Genes were predicted from the MAGs with Prokka v1.11 (Seemann, 2014).

A Bacterial MAG phylogenomic tree based on 120 concatenated marker genes from GTDB-tk (Chaumeil et al., 2020) was constructed by FastTree v2.1.11 (Price et al., 2010). The tree was visualized with Interactive Tree of Life (Letunic and Bork, 2024).

Eukaryotic bins were recovered along with the prokaryotic bins in our assembly and binning workflow. Large (>5Mbp) eukaryote-like bins, often initially annotated as archaea by checkm, were assessed with 83 eukaryotic single copy genes for completeness and contamination with anvi’o (Eren et al., 2021). The good quality bins (>=50% completeness and <=11% contamination) were dereplicated with dRep. Pairwise ANI values were checked to further dereplicate the bins (cutoff 95%). The closest references of the bins were first determined by finding ANI matches to reference genomes using FastANI (Jain et al., 2018). If an ANI match was not available, 5S rRNA and anvi’o protist marker genes predicted from the bins were searched against NCBI Nucleotide/Refseq databases. Our workflow resulted in 11 unique good quality eukaryotic bins.

### Hydrological and flow cytometry data

Methods for processing and quantifying environmental parameters can be found in the BCO-DMO repository (Project ID: #935794 and #935786). Flow cytometry data for each sampling was collected by the Paerl Lab at North Carolina State University (Raleigh, North Carolina, USA).

## RESULTS and DISCUSSION

### Environmental variability across sampling sites

From August 2021 to August 2022, we collected surface-water samples (∼1 m depth) at three stations spanning the estuary-to-ocean continuum: NRE180 in the lower Neuse River Estuary (more riverine western, up-estuary), PS5 in Pamlico Sound (higher saline, eastern estuary), and OB1 in Onslow Bay (Fig. 1A and 1B). NRE180 and PS5 are part of a long-term monitoring effort (MODMON). These sites had similar temperatures and dissolved-oxygen concentrations (SI Fig. 1), yet differed markedly in hydrologic regime, as shown by a pronounced salinity gradient from the estuary to coastal waters (Fig. 1C).

Mean chlorophyll-*a* concentrations indicated that NRE180 and PS5 were more productive than OB1, averaging 7.2 and 6.7 µg Chl *a* L⁻¹, respectively, compared with 0.9 µg Chl *a* L⁻¹ offshore (Fig. 1D). NRE180 had elevated POC, PN, DOC, pico- and nanoeukaryote abundances, and prokaryotic cell densities, whereas PS5 had greater numbers of >10 µm eukaryotes and Synechococcus-like cells (SI Fig. 1; SI Table 1). OB1 had the lowest eukaryotic and prokaryotic cell counts, consistent with its more oligotrophic character.

### Microbial community compositions

Over the year-long survey we generated 17 quantitative metagenomes, six each from the Neuse River Estuary (NRE180) and Pamlico Sound (PS5) and five from the offshore station in Onslow Bay (OB1).

Read-level taxonomic profiling against NCBI RefSeq (v223) showed that the metagenomes were overwhelmingly bacterial. Across the 17 libraries, on average >90% of quality-filtered reads mapped to Bacteria; the remainder affiliated with Eukaryota (∼5%), viruses (3%), and Archaea (0.3%) (SI Fig. 2). Composition was notably temporally consistent in the estuarine (NRE180) and sound (PS5) datasets, whereas the offshore OB1 libraries harbored a notable enrichment of eukaryotic reads accompanied by a proportional decrease in bacterial reads in November (O2) and January (O3) samples. This is consistent with the flow cytometry data from the offshore station from these two samples, which had relatively low bacterial cell counts and relatively high picoeukaryote (<2 µm) abundances (SI Table 1).

**Figure 2.**
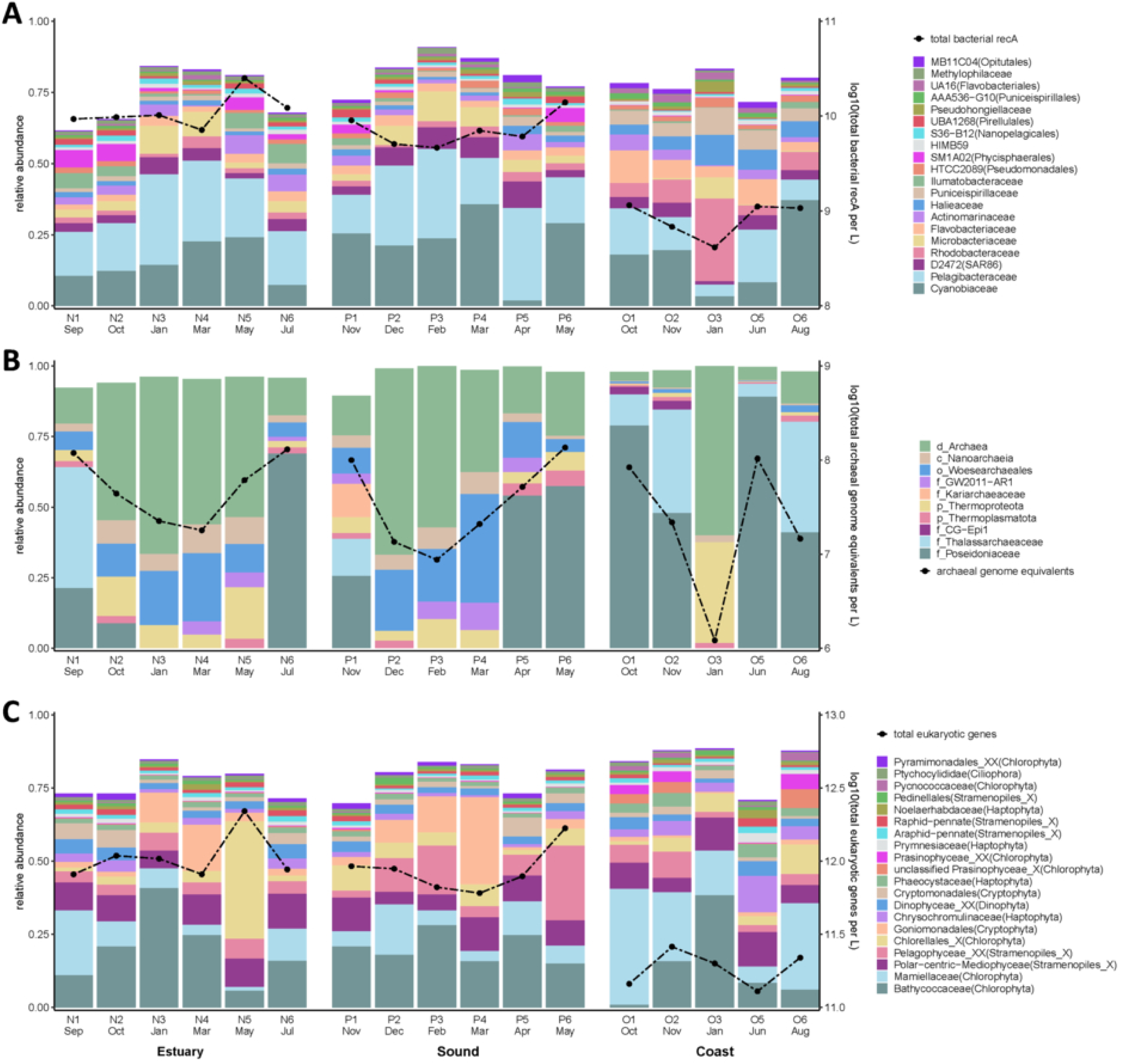
Spatiotemporal dynamics of microbial community composition across the estuarine to coastal ocean continuum. (A) Bacterial community structure based on abundance of *recA* reads classified at the family level, (B) archaeal community composition based on abundance of reads annotated with SingleM, and (C) eukaryotic community composition based on abundance of reads annotated with PhyloDB. In all panels, stacked bars represent the relative abundance of dominant taxonomic groups for each sampling event at the Estuary (NRE180), Sound (PS5), and Coast (OB1) stations. The secondary y-axis and superimposed black dashed lines represent absolute volumetric abundances: total bacterial *recA* genes L^-1^ (A), total archaeal genome equivalents L^-1^ (B), and total eukaryotic genes L^-1^ (C), determined through internal standard normalization.

### Bacteria metagenome-derived abundances

To convert metagenomic read counts into volumetric cell abundances, we targeted the universal, single-copy bacterial gene *recA* (Landa et al., 2019). Reads mapping to *recA* were taxonomically classified with the genome taxonomy database (GTDB), and their counts were normalized with the internal-standard genomes to yield volumetric gene abundances (genomes L^-1^). Each bacterial genome carries a single *recA* locus, so these values correspond to genome equivalents assuming on average one genome per cell. (Fig. 2A).

Across the estuarine (NRE180) and sound (PS5) stations, total *recA*-derived cell densities averaged 1 × 10¹⁰ cells L⁻¹ (Fig. 2A). The offshore station OB1 was an order of magnitude less populous, averaging 9 × 10⁸ cells L⁻¹, mirroring its lower productivity and phytoplankton biomass (Fig. 1D). Bacterial abundances at all three locations declined during winter, although the minima did not coincide precisely with the temperature minimum (Fig. 2A; SI Fig. 1), indicating that additional factors, such as substrate availability or grazer pressure (Lima-Mendez et al., 2015), modulate seasonal biomass fluctuations.

Across all samples, *recA*-derived profiles were dominated by a core set of bacterial families, with Cyanobiaceae and Pelagibacteraceae almost invariably ranking first and second in relative abundance (Fig. 2A) and averaging 1.4 × 10^9^ and 1.5 × 10^9^ cells L⁻¹, respectively (SI Table 3). Secondary contributors (typical abundances 2.5 × 10^8^ cells L⁻¹) included the SAR86 family D2472, Rhodobacteraceae, Microbacteriaceae, and Flavobacteriaceae. The estuarine stations NRE180 and PS5 shared broadly similar compositions and were particularly enriched in Microbacteriaceae and D2472, whereas the offshore site (OB1) displayed higher proportions of Rhodobacteraceae, Flavobacteriaceae, Halieaceae, and *Candidatus* Puniceispirillaceae (SAR116).

Cyanobiaceae and Pelagibacteraceae generally remained the two most abundant families, with two notable exceptions: the April Pamlico Sound sample (PS5) and the January OB1 offshore sample (O3). Seasonal signals were clearest at estuarine stations, where cooler months were characterized by enrichment of Cyanobiaceae, Pelagibacteraceae, and Microbacteriaceae. Seasonal interpretation at OB1 is complicated by a large temporal gap in sampling between January and June. Nevertheless, the January offshore sample (O3) stood out as compositionally distinct, exhibiting increased Rhodobacteraceae, Microbacteriaceae, and Halieaceae, with the relative decrease of Cyanobiaceae, Pelagibacteraceae, and Flavobacteriaceae. The offshore site composition exhibited a predominantly marine signal. Its community structure was broadly similar to the nearby Pivers Island Coastal Observatory (PICO) time-series (Ward et al., 2017) but with a conspicuously lower representation of Pelagibacteraceae (Fig. 2A). Indeed, Pelagibacteraceae constituted a smaller fraction of the offshore bacterial assemblage than at the more nutrient-rich stations estuarine and sound sites. (Morris et al., 2002; Brown et al., 2012) highlighting its ability to thrive in various environments (Campbell and Kirchman, 2013; Campbell et al., 2022).

### Archaea metagenome-derived abundances

Archaeal reads classified with SingleM and normalized with internal standards showed that archaeal genome equivalents were one to two orders of magnitude lower than bacterial abundances but varied seasonally, declining to ∼10⁷ L⁻¹ in cooler months and increasing to ∼10⁸ L⁻¹ during summer (Fig. 2B). Warm-season archaeal communities were dominated by Marine Group II families Poseidoniaceae and Thalassarchaeaceae, consistent with their prevalence in euphotic coastal waters, whereas cooler months contained more diverse but lower-abundance archaeal assemblages, including Woesearchaeales at the estuarine and sound stations and Thermoproteota in the January offshore sample. Thaumarchaeota and other ammonia-oxidizing lineages were minor components, potentially reflecting the low ammonia concentrations observed across most samples. Overall, archaeal abundances and composition showed a strong seasonal signal but represented a small fraction of the total microbial community.

### Eukaryotic Metagenome-derived abundances

Eukaryotic reads were taxonomically assigned with PhyloDB and scaled with internal standard conversion factors to generate semi-quantitative gene abundance estimates (Fig. 2C). Because the 0.2–3 µm size fraction selectively captures pico- and small nanoeukaryotes, and because total eukaryotic gene counts cannot be directly converted to genome equivalents, these values should be interpreted as indicators of relative cell abundance across samples. Eukaryotic gene counts were generally higher at the estuarine and sound stations than offshore, with spring peaks at both APES sites. Chlorophytes, especially Bathycoccaceae, Mamiellaceae, and Chlorellales, accounted for much of the annotated eukaryotic signal (SI Fig. 3), consistent with prior evidence that small chlorophytes are important members of the pico- and nanoplankton (Vaulot et al., 2008; Gaulke et al., 2010; Gong et al., 2020).

**Figure 3.**
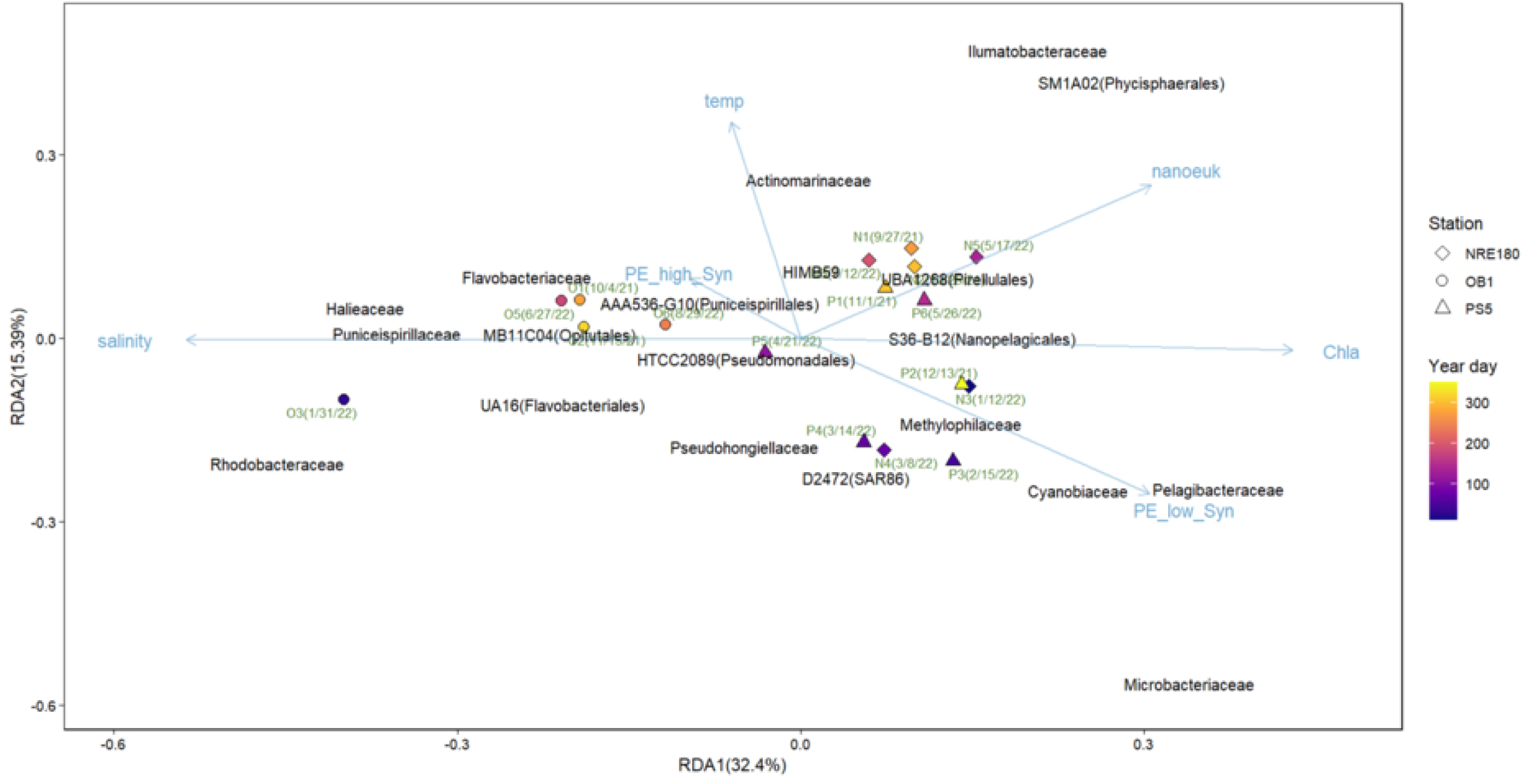
Redundancy analysis (RDA) of microbial community composition and environmental variables across the study transect. The triplot illustrates relationships between sampling events (points), bacterial families (labels), and environmental drivers (blue arrows). Community composition is based on the 20 most abundant bacterial families identified via *recA* read-based survey. RDA1 (32.4%) and RDA2 (15.3%) collectively account for 47.7% of the total variance in community structure. Points are shaped by station (NRE180 = diamonds, PS5 = triangles, OB1 = circles) and colored according to the sampling day of the year. Environmental vectors include surface temperature (temp), salinity, chlorophyll-a (Chla), nanoeukaryote abundance (nanoeuk), and two cytometric clusters of *Synechococcus* defined by high (PE_high_Syn) or low (PE_low_Syn) phycoerythrin fluorescence.

### Community shifts and environmental drivers

Redundancy analysis (RDA) of the 20 most abundant recA-defined bacterial families explained 48% of community variation across the first two axes, revealing a primary spatial separation between offshore OB1 and the estuarine NRE180 and PS5 sites along the salinity-associated RDA1 axis, and a secondary seasonal gradient along RDA2 (Fig. 3). Offshore communities were associated with higher salinity and greater relative abundances of Halieaceae, Flavobacteriaceae, Ca. Puniceispirillaceae, UA16, and MB11C04, while estuarine and sound communities showed stronger seasonal structure, with summer-to-autumn samples associated with warmer temperatures, phytoplankton-related variables, and increased Ilumatobacteraceae, Actinomarinaceae, and SM1A02. Winter and spring samples were characterized by higher relative abundances of Pelagibacteraceae, Cyanobiaceae, and Microbacteriaceae. Overall, the ordination indicates that salinity structured the major estuarine-to-coastal biogeographic gradient, while temperature and correlated phytoplankton/cyanobacterial variables contributed to seasonal community succession within APES, consistent with previous studies showing strong effects of salinity and temperature on coastal microbial communities (Fortunato et al., 2012; Chow et al., 2013; Ward et al., 2017; Sanchez-Gallego et al., 2025).

### MAG recovery and quantitative coverage of microbial populations

To resolve community dynamics at the population level, we reconstructed MAGs from each of the 17 libraries, generating 415 MAGs: 386 bacterial, 18 archaeal, and 11 eukaryotic. Prokaryotic MAGs were highly complete overall, with a median completeness of 83% and median genome size of 2.1 Mb, spanning 18 phyla and more than 100 families, including dominant groups identified in the read-based survey such as Cyanobiaceae, Pelagibacteraceae, and *Ca.* Puniceispirillaceae (Fig. 4; SI Table 4).

**Figure 4.**
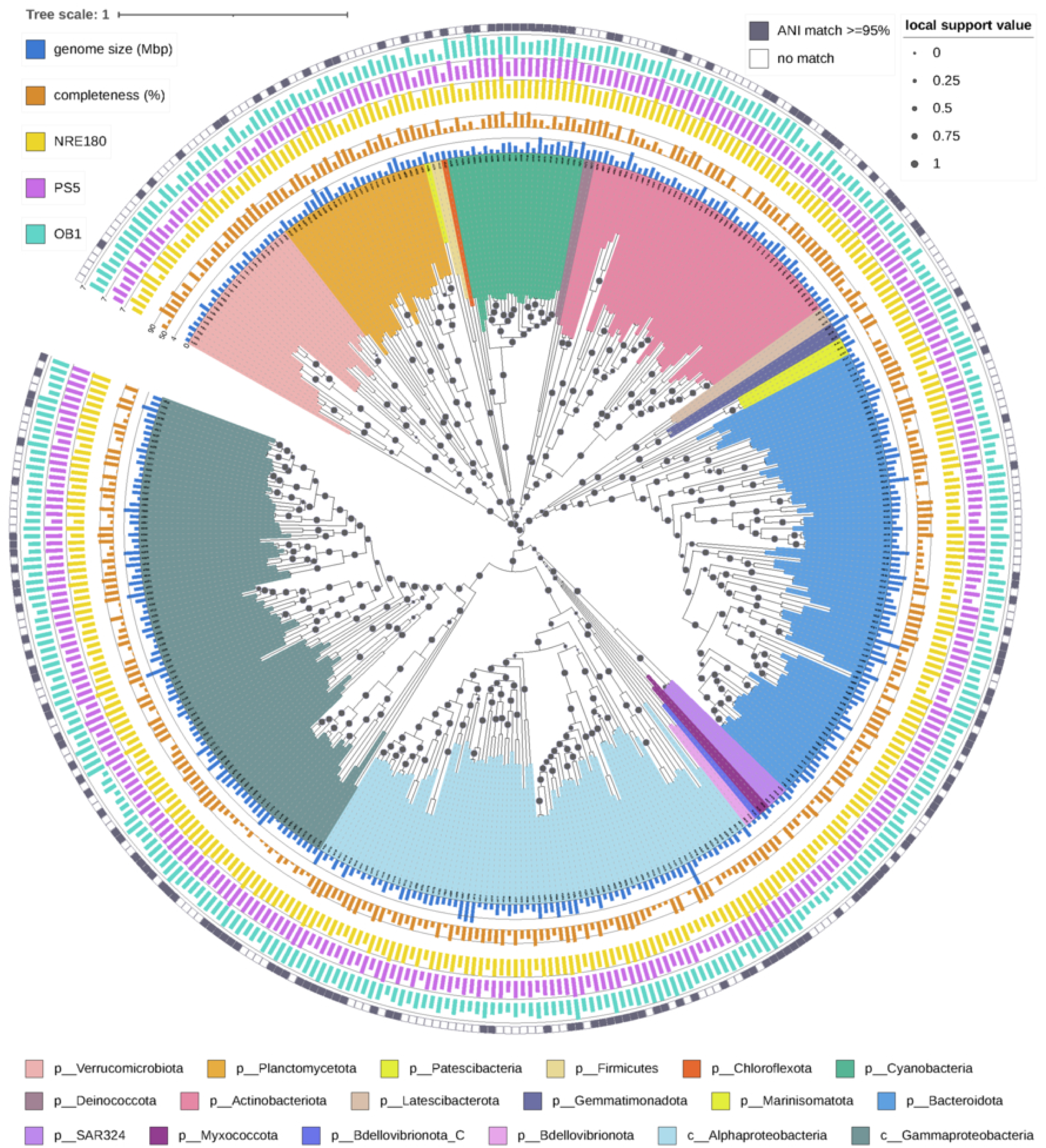
Phylogenomic diversity and characteristics of 386 bacterial metagenome-assembled genomes (MAGs) recovered from the study transect. The circular maximum-likelihood tree is based on a concatenated alignment of 120 single-copy marker genes, with local support values at nodes indicated by grey circles. Internal leaf colors represent taxonomic classification at the phylum or class level according to the Genome Taxonomy Database (GTDB). Successive outer rings display, from inside to out: genome size in Mbp (blue bars), genome completeness (orange bars), and log-transformed average absolute abundances (genome equivalents L^-1^) at the estuarine (NRE180; yellow), sound (PS5; purple), and offshore (OB1; teal) stations. The outermost ring consists of colored squares indicating MAGs with a ≥ 95% Average Nucleotide Identity (ANI) match to existing reference genomes.

Internal standard normalization converted MAG coverage values to absolute genome equivalents, showing that the MAGs captured, on average, 52% of bacterial, 20% of archaeal, and ∼50% of picoeukaryote abundances across samples (Fig. 5; SI Fig. 6 and 7). Among bacterial MAGs, 166 shared ≥95% ANI with GTDB reference genomes, whereas 220 lacked a close reference match, indicating that many abundant coastal and estuarine populations remain poorly represented in genome databases. Several novel populations reached 10⁷–10⁸ genome equivalents L⁻¹, demonstrating that the recovered MAGs represent a substantial and quantitatively important fraction of the community rather than rare background taxa.

**Figure 5.**
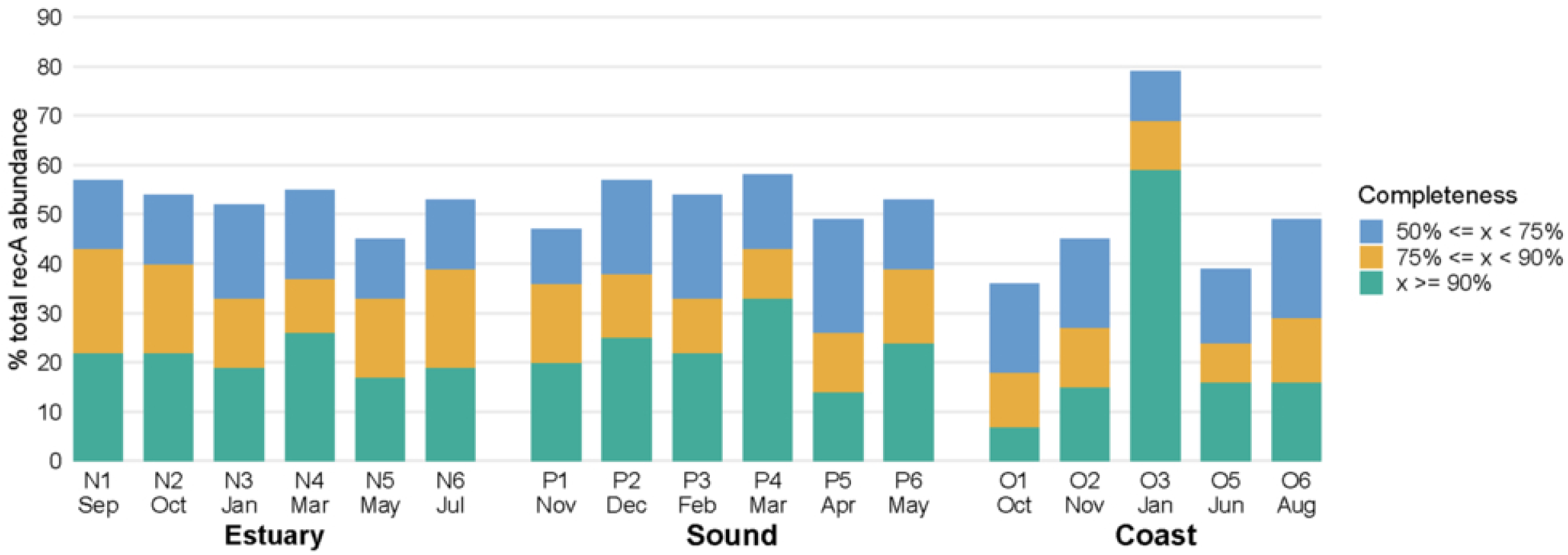
Proportion of total bacterial abundance represented by metagenome-assembled genomes (MAGs) across the estuary-to-ocean continuum. Stacked bar plots indicate the cumulative percentage of total bacterial genome equivalents captured by the 386 bacterial MAGs in each of the 17 quantitative metagenomes. Values are calculated by comparing the summed absolute abundances of MAGs to total *recA*-derived bacterial genome equivalents. Bars are categorized by MAG completeness thresholds: ≥90 (green), ≥ 75% (yellow), and ≥ 50% (blue). Categories are nested, with higher completeness groups representing subsets of the lower completeness tiers. Sample codes on the x-axis correspond to the dates and stations (Estuary: NRE180, Sound: PS5, Coast: OB1) detailed in Figure 1B.

**Figure 6.**
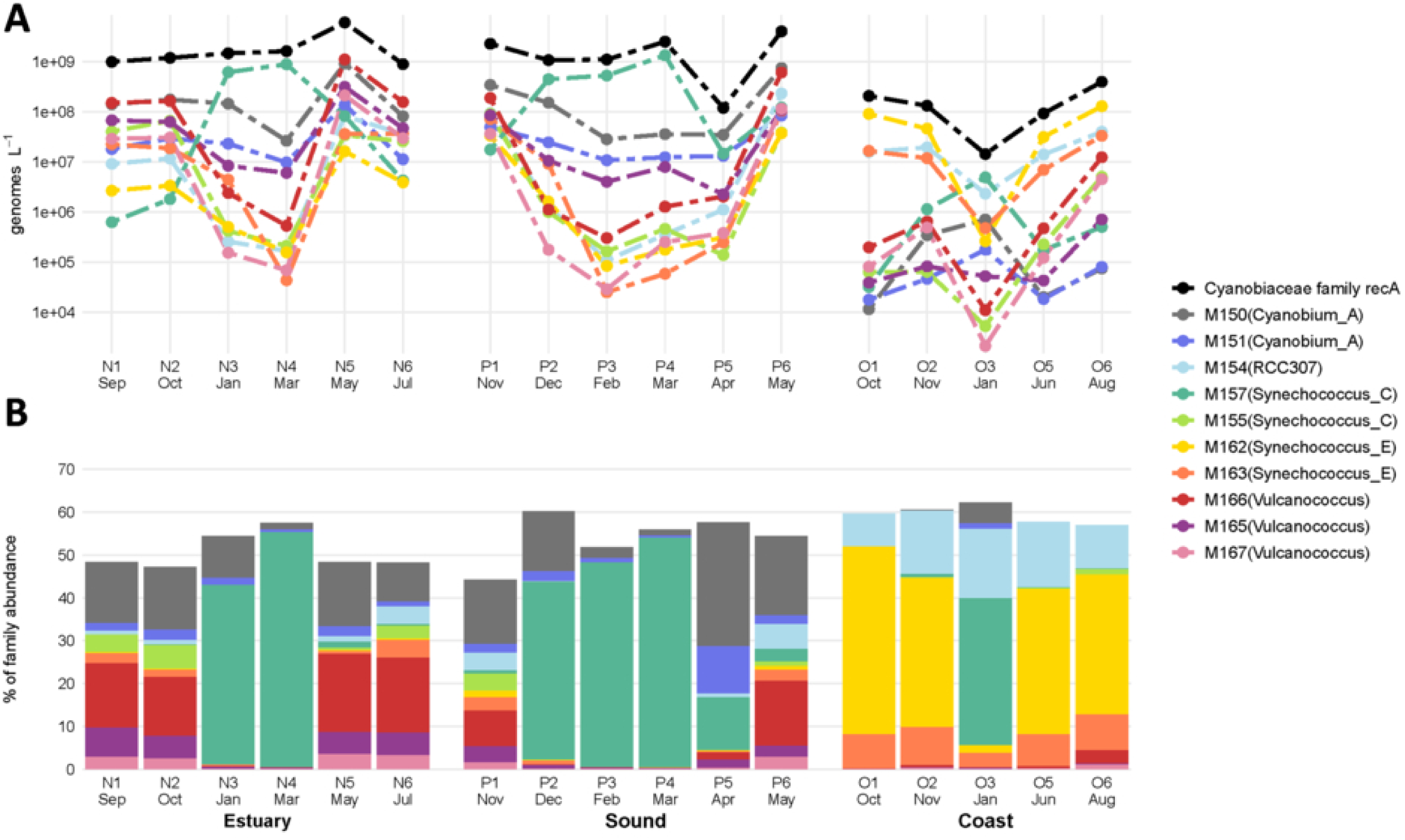
Absolute and relative abundances of the top 10 most abundant Cyanobiaceae metagenome-assembled genomes (MAGs). (A) Volumetric absolute abundances (genomes L^-1^) of the 10 most abundant Cyanobiaceae MAGs across the 17 sampling events. The black dashed line represents the total Cyanobiaceae family absolute abundance derived from Cyanobiaceae *recA* gene counts. (B) Relative abundance of each MAG as a percentage of the total Cyanobiaceae *recA* abundance. MAGs are identified by their ID followed by the GTDB-tk assigned genus in parentheses. Data are organized by sampling station: Estuary (NRE180), Sound (PS5), and Coast (OB1) with sample codes corresponding to dates detailed in Figure 1B.

**Figure 7.**
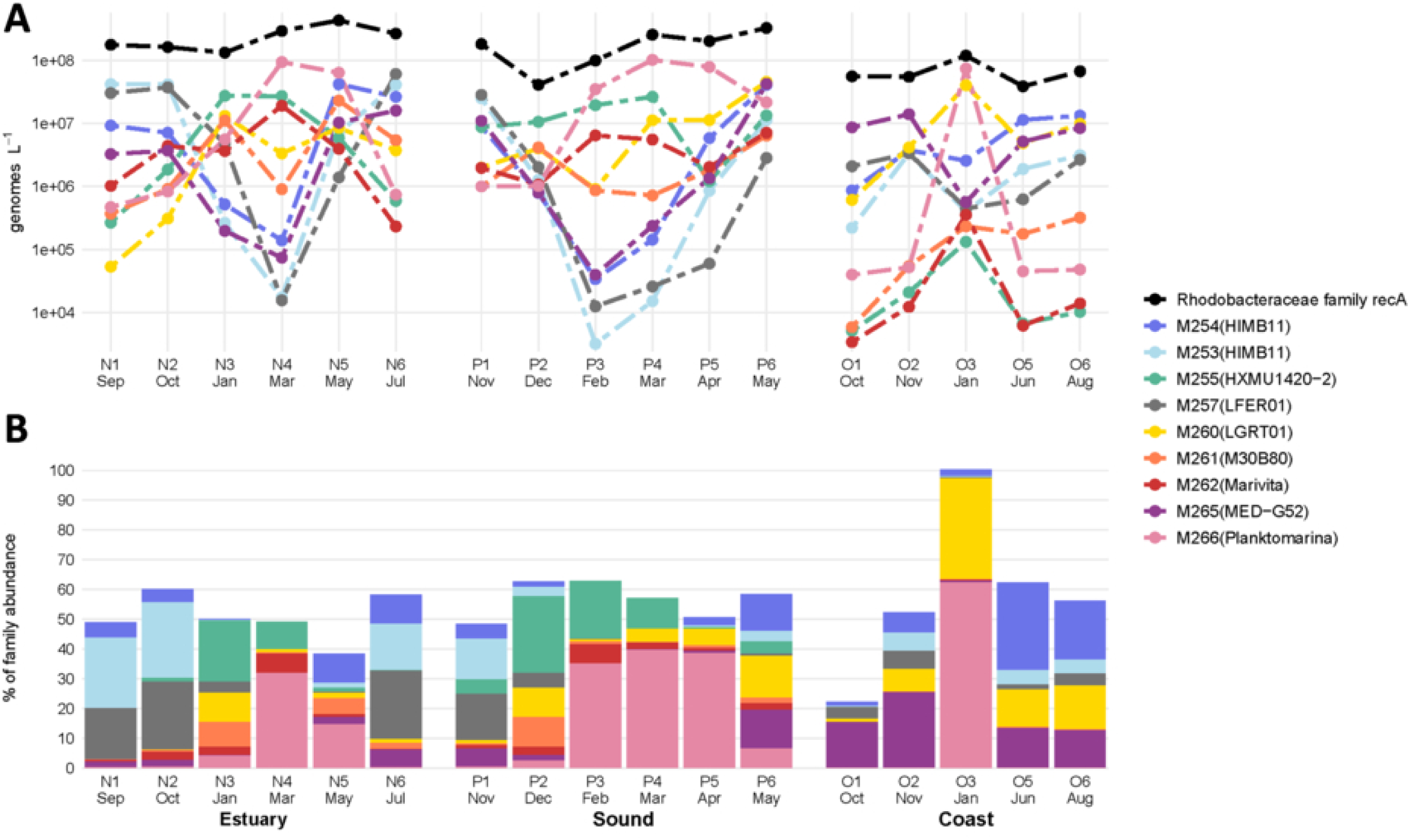
Absolute and relative abundances of the top nine most abundant *Rhodobacteraceae* metagenome-assembled genomes (MAGs). (A) Volumetric absolute abundances (genomes L-1) of the primary *Rhodobacteraceae* MAGs across the 17 sampling events. The black dashed line indicates the total *Rhodobacteraceae* family absolute abundance determined from *recA* gene counts. (B) Relative abundance of each MAG expressed as a percentage of the total *Rhodobacteraceae recA* abundance. MAG IDs are listed with their GTDB-tk assigned genus in parentheses. Data are organized by sampling station: Estuary (NRE180), Sound (PS5), and Coast (OB1) with sample codes corresponding to dates detailed in Figure 1B.

### Bacterial MAG Recovery and Community Coverage

The bacterial MAG collection captured a substantial and seasonally responsive fraction of the community. A phylogenomic reconstruction based on the concatenated alignment of 120 single-copy marker genes used byGTDB-Tk placed the bacterial MAG collection into a well-resolved tree that highlights both its breadth and novelty (Fig. 4). The 386 genomes span 17 bacterial phyla, with the richest representation in the classes Alphaproteobacteria and Gammaproteobacteria, and the phyla Bacteroidota, Actinobacteria, Cyanobacteria, Planctomycetota, and Verucomicrobiota, a pattern that mirrors the dominant families identified in the *recA* read-based survey (Fig. 2A).

MAG representation varied modestly among estuarine and sound samples, ranging from 45–58% of total bacterial genome equivalents (Fig. 5), and peaked at 79% in the winter offshore sample O3, which was enriched in *Planktomarina* MAG M266 (SI Fig. 5). Applying a stricter ≥90% completeness threshold reduced mean bacterial coverage to 22% (Fig. 5), underscoring the difficulty of recovering near-complete genomes for abundant taxa in complex estuarine and coastal microbiomes.

### Archaeal MAGs

Genome reconstruction yielded 18 archaeal MAGs, all assigned by GTDB-Tk to Marine Group II (MGII) families Poseidoniaceae and Thalassarchaeaceae (SI Table 4). This exclusive recovery of MGII lineages mirrors the dominance of these taxa in the archaeal non-assembled analysis (Fig. 2B), underscoring their numerical importance in surface waters along the estuary-to-ocean transect.

When the volumetrically normalized abundances of these archaeal MAGs were compared with the archaeal genome-equivalent counts derived from singleM (SI Fig. 6), the fraction of the community captured by the archaeal MAG catalogue ranged from 0% to 57% across samples, with a mean of ∼20%. The highest percentages coincided with summer offshore OB1 (O5) and estuarine NRE180 (N1) samples, where MGII abundances peaked. The winter and early-spring libraries, characterized by more taxonomically diverse but low-density archaeal assemblages, contributed little or nothing to the MAG total.

The variable recovery efficiency suggests that most non-MGII archaeal populations were present at coverages too low to assemble into genomes with current short-read methods. Thus, while the MGII MAGs provide genomic insight into the dominant archaea which are roughly four-fifths of the archaeal cells, nearly all taxonomic diversity outside MGII remain unresolved at the genome level, highlighting an important frontier for deeper or long-read sequencing approaches.

### Eukaryotic MAGs

Eukaryotic genome reconstruction is inherently more challenging than prokaryotic assembly because of larger genome sizes, intron density, and heterozygosity (Alexander et al., 2023). Even so, our workflow yielded 11 high-quality eukaryotic MAGs (≥50% completeness, ≤11% contamination; SI Table 5). Closest relatives, established by ≥74.9% ANI matches or BLAST matches of 5S rRNA and conserved eukaryotic marker genes (see Methods), place these genomes within the dominant picophytoplankton lineages *Micromonas*, *Bathycoccus*, and *Picochlorum*, all chlorophytes with compact genomes well suited to the 0.2-3 µm size fraction captured on our filters. Taxa represented by the MAGs were also found to be abundant in our non-assembled analysis (Fig. 2C).

To gauge their quantitative importance, we compared volumetrically normalized eukaryotic MAG coverages with flow-cytometric counts of picoeukaryotic algae (<2µm diameter) (SI Fig. 7). On average, the 11 MAGs account for ∼50% of the counted picoeukaryotes, confirming that they represent a substantial share of the phototrophic nano- and picoplankton community. The correspondence varies among samples, from a low of 12% (N1) to a high of 129% of picoeukaryote flow cytometry counts (O1). Overestimation at offshore O1 likely reflects the combined effects of lower overall eukaryotic read depth and the inherent uncertainty of converting short-read coverage to absolute genome equivalents when coverage is sparse.

Recent advances in the reconstruction of eukaryotic MAGs have begun to illuminate the poorly resolved branches of the eukaryotic tree of life and the functional repertoires of environmentally relevant eukaryotic organisms (Delmont et al., 2022; Alexander et al., 2023, Bei et al., 2025). While chlorophyte taxa such as Chlorellales and Mamiellales are known to constitute a significant portion of the eukaryotic community in the Neuse River Estuary (Gong et al., 2020), several of the MAGs we recovered are distantly related to the available reference genomes, suggesting they may represent locally adapted chlorophyte populations.

### Cyanobiaceae Genomic Insights and Pigment-Based Signatures

The family Cyanobiaceae, which encompasses the globally abundant genera *Synechococcus*, *Prochlorococcus*, and *Cyanobium* (Parks et al., 2022), was the most prominent cyanobacterial lineage in our data set (Fig. 2A). We reconstructed 21 Cyanobiaceae MAGs that collectively span the dominant *Synechococcus* clades in the region (Fig. 4; SI Table 6). Seven have no ≥95% ANI match in public databases, and only two of the remaining 14 have ANI matches exceeding 99%, indicating substantial genomic novelty among the MAGs. Closest references were mostly from the northwestern Atlantic margin, Chesapeake Bay, Woods Hole, and the Gulf of Mexico, indicating broad regional representation among related picocyanobacterial populations (SI Table 6).

The ten most abundant Cyanobiaceae MAGs accounted for ∼55% of Cyanobiaceae genome equivalents in each sample (Fig. 6B) and displayed pronounced, site-specific seasonal patterns. In estuarine NRE180 and sound PS5 samples, winter and early-spring communities were dominated by M157 (Synechococcus_C), whereas summer–autumn assemblages shifted toward M166 (*Vulcanococcus*) and M150 (*Cyanobium*_A).

At the offshore station OB1, M162 (Synechococcus_E) prevailed throughout the year except in the January sample (O3), when the abundances of M157, M150, M151 (Cyanobium_A) and M165 (Vulcanococcus) all increased despite having lower absolute abundances than observed in the estuary and sound sites (Fig. 6A). The collective behavior of these MAGs therefore marks a clear transition from winter–spring communities dominated by a single *Synechococcus*_C population to diverse summer–autumn assemblages that differ between estuarine and coastal waters, reflecting niche adaptation and mechanisms for population persistence (e.g. populations are not readily flushed out of the system) along the salinity–turbidity gradient.

Distributions of the top Cyanobiaceae MAGs demonstrate that even though these cyanobacteria populations were always present to some degree, very different compositions were found at the estuary compared to in the coastal water, indicating adaptations to either coastal or estuarine environments. Our analyses found Cyanobiaceae MAGs correlated with flow cytometry counts of distinctly pigmented *Synechococcus*-like cells (SI Fig. 8). M157 (*Synechococcus*_C) and M162 (*Synechococcus*_E) track PE-high and PE-low *Synechococcus* counts, respectively. While both are likely PE-rich strains that tend to dominate in less turbid coastal ocean water and were abundant in the coastal OB1 site, their dominance at OB1 switched in the winter, suggesting seasonal niche partitioning that might be associated with PE-high and PE-low pigment profiles (Lantoine and Neveux, 1997; Stomp et al., 2007; Haverkamp et al., 2009; Sanfilippo et al., 2019). In contrast, M155 (*Synechococcus*_C), M150 (*Cyanobium*_A), M151 (*Cyanobium*_A), and M165 (*Vulcanococcus*) correlate with PC-high *Synechococcus* cells that preferentially inhabit the turbid, nutrient-rich estuary (SI Fig. 8) (Stomp et al., 2007).

Our results demonstrate that genome-resolved time series can reveal fine-scale cyanobacterial population dynamics that are obscured by pigment-based or bulk community measurements alone. Although flow cytometry distinguishes broad PE- and PC-containing groups, MAG-based abundances resolved seasonal turnover and shifts in dominance among closely related lineages within these categories. These patterns are consistent with niche partitioning linked to light quality and turbidity, which are known to structure *Synechococcus* pigment ecotypes (Stomp et al., 2007; Haverkamp et al., 2009), and with prior work connecting pigment diversity to genomic variation and flow cytometry-defined pigment classes (Six et al., 2007; Scanlan et al., 2009; Li et al., 2024). While direct links between MAG features and in situ pigment phenotypes remain uncertain, tracking individual populations through time provides new insight into how closely related cyanobacterial lineages partition niches across estuarine–coastal gradients.

### Rhodobacteraceae MAGS and their Eukaryotic associations

Rhodobacteraceae are metabolically versatile heterotrophs that thrive in estuarine and coastal waters worldwide (Buchan et al., 2005), and they were likewise persistent, and often abundant in our samples (Fig. 7A). We reconstructed 19 Rhodobacteraceae MAGs, 13 of which share ≥95% ANI with previously published genomes. In contrast to the cyanobacterial references that are dominated by isolates, the closest relatives of most of our Rhodobacteraceae MAGs are themselves environmental MAGs collected from disparate ocean basins (SI Table 7).

The nine most prolific MAGs together accounted for 55% of Rhodobacteraceae genome equivalents on average (Fig. 7B), yet they exhibited pronounced spatial and seasonal turnover. Four populations, M253 and M254 from HIMB11, M265 (MED-G52), and M257 (LFER01), declined sharply during winter and early spring at estuarine NRE180 and sound PS5 and were likewise suppressed in the January offshore sample O3. Conversely, the other MAGs peaked during the colder months, but their timing and magnitude varied among stations.

The most striking Rhodobacteraceae MAG distribution was M266 (*Planktomarina;* 99% ANI match to *Planktomarina temperata* RCA23), which became the prominent Rhodobacteraceae MAG in late winter and spring at the estuarine and sound site. At the offshore site, it comprised ∼60 % of family abundance and 18 % of all bacterial cells in sample O3 (Fig. 7B; SI Fig. 5). Despite its streamlined genome of 2.5 Mbp, smaller than the 3.3 Mb RCA23 genome, M266 evidently competes successfully under winter conditions along the shelf. Other MAGs showed clear habitat preferences: M253, M257, M255 (HXMU1420-2) were most common in the estuary and sound, whereas M265 and M260 (LGRT01) were enriched offshore. Together, the distributions of the Rhodobacteraceae MAGs indicate niche partitioning both in space and season.

Many Rhodobacteraceae lineages are associated with phytoplankton (Buchan et al., 2005; Kieft et al., 2021; Isaac et al., 2024), so we compared MAG abundances with flow-cytometric phytoplankton measurements and with the 11 eukaryotic MAGs recovered from the same samples. Only modest correlations emerged with FCM phytoplankton counts. For instance, M255 (HXMU1420) tracked *Synechococcus* and M261 (M30B80) tracked picoeukaryotic phytoplankton abundance (SI Fig. 9). Stronger and highly specific relationships appeared between individual Rhodobacteraceae and eukaryotic MAGs (SI Fig. 10). Notably, the abundant *Planktomarina* (M266) co-varied with a *Micromonas* MAG (M415), suggesting that population-level interactions, rather than aggregate phytoplankton biomass, structure these bacterium–alga partnerships.

Such population-level associations are consistent with growing evidence that phytoplankton–bacteria interactions can be specific rather than generalized, particularly among Rhodobacterales and phytoplankton (Grossart et al., 2005; Buchan et al., 2014; Lima-Mendez et al., 2015). These interactions are often mediated by exchanges of organic carbon, vitamins, nitrogen, and sulfur compounds, providing a plausible basis for the observed co-variation (Miller and Belas, 2004; Han et al., 2021). Although these correlations do not demonstrate direct interaction or causality, the weak correspondence with bulk phytoplankton abundance but strong MAG-level correlations suggest that microbial associations in this system are structured at finer taxonomic resolution than traditional measurements capture.

By quantifying individual genome-resolved populations across the estuarine-to-coastal continuum, this study shows how absolute abundance measurements can transform metagenomic surveys from descriptions of community composition into numerical frameworks for linking microbial population dynamics to coastal ecosystem function.

## Acknowledgements

We thank Jeremy Braddy and Amy Bartenfelder for assistance with water collections. We thank Dr. Hans Paerls and the MODMON team for analyzing and providing hydrological data. We thank Garrett Sharpe, Melanie Cohn, David Malcolm, and Savannah Curtis for assisting in sampling trips. This work was financially supported by the United States National Science Foundation Division of Ocean Sciences grants OCE-1850692 and OCE-2505930 to S.M.G.; and OCE-2049388 and OCE-2416286 to R.W.P.

## Data availability

Raw sequence reads are available in the NCBI SRA under BioProject no. PRJNA1157453.

## REFERENCES

Alexander, H., Hu, S. K., Krinos, A. I., Pachiadaki, M., Tully, B. J., Neely, C. J., et al. (2023). Eukaryotic genomes from a global metagenomic data set illuminate trophic modes and biogeography of ocean plankton. mBio 14, e01676–23. doi: 10.1128/mbio.01676-23

Azam, F., and Malfatti, F. (2007). Microbial structuring of marine ecosystems. Nat Rev Microbiol 5, 782–791. doi: 10.1038/nrmicro1747

Bei Q, Williams NL, Furtado LE, Blasi DD, Williams J, Brotas V, Tarran G, Rees AP, Bowler C, Fuhrman JA. Quantitative metagenomics for marine prokaryotes and photosynthetic eukaryotes. ISME communications. 2025 Jan;5(1):ycaf131.

Bolger, A. M., Lohse, M., and Usadel, B. (2014). Trimmomatic: a flexible trimmer for Illumina sequence data. Bioinformatics 30, 2114–2120. doi: 10.1093/bioinformatics/btu170

Brown, M. V., Lauro, F. M., DeMaere, M. Z., Muir, L., Wilkins, D., Thomas, T., et al. (2012). Global biogeography of SAR11 marine bacteria. Molecular Systems Biology 8, 595. doi: 10.1038/msb.2012.28

Buchan, A., González, J. M., and Moran, M. A. (2005). Overview of the Marine Roseobacter Lineage. Applied and Environmental Microbiology 71, 5665–5677. doi: 10.1128/AEM.71.10.5665-5677.2005

Buchan, A., LeCleir, G. R., Gulvik, C. A., and González, J. M. (2014). Master recyclers: features and functions of bacteria associated with phytoplankton blooms. Nature Reviews Microbiology 12, 686–698. doi: 10.1038/nrmicro3326

Buchfink, B., Xie, C., and Huson, D. H. (2015). Fast and sensitive protein alignment using DIAMOND. Nat Methods 12, 59–60. doi: 10.1038/nmeth.3176

Buzzelli, C., Luettich, R., Paerl, H., Fear, J., Fleming, J., Twomey, L., et al. (2002). Neuse River Estuary Modeling and Monitoring Project Phase 2: Monitoring of Hydrography and Water Quality, Circulation, Phytoplankton Physiology and Sediment-Water Coupling. Available at: https://www.semanticscholar.org/paper/Neuse-River-Estuary-Modeling-and-Monitoring-Project-Buzzelli-Luettich/d2e1f92158c7bb7db1744469e2ef0e0ea256ec82 (Accessed June 6, 2025).

Campbell, B. J., and Kirchman, D. L. (2013). Bacterial diversity, community structure and potential growth rates along an estuarine salinity gradient. The ISME Journal 7, 210–220. doi: 10.1038/ismej.2012.93

Campbell, B. J., Lim, S. J., and Kirchman, D. L. (2022). Controls of SAR11 subclade abundance, diversity, and growth in two Mid-Atlantic estuaries. 2022.05.04.490708. *Biorxiv* doi: 10.1101/2022.05.04.490708

Chaumeil, P.-A., Mussig, A. J., Hugenholtz, P., and Parks, D. H. (2020). GTDB-Tk: a toolkit to classify genomes with the Genome Taxonomy Database. Bioinformatics 36, 1925–1927. doi: 10.1093/bioinformatics/btz848

Chow, C.-E. T., Sachdeva, R., Cram, J. A., Steele, J. A., Needham, D. M., Patel, A., et al. (2013). Temporal variability and coherence of euphotic zone bacterial communities over a decade in the Southern California Bight. ISME J 7, 2259–2273. doi: 10.1038/ismej.2013.122

Delmont, T. O., Gaia, M., Hinsinger, D. D., Frémont, P., Vanni, C., Fernandez-Guerra, A., et al. (2022). Functional repertoire convergence of distantly related eukaryotic plankton lineages abundant in the sunlit ocean. Cell Genomics 2. doi: 10.1016/j.xgen.2022.100123

Eren, A. M., Kiefl, E., Shaiber, A., Veseli, I., Miller, S. E., Schechter, M. S., et al. (2021). Community-led, integrated, reproducible multi-omics with anvi’o. Nat Microbiol 6, 3–6. doi: 10.1038/s41564-020-00834-3

Fortunato, C. S., Herfort, L., Zuber, P., Baptista, A. M., and Crump, B. C. (2012). Spatial variability overwhelms seasonal patterns in bacterioplankton communities across a river to ocean gradient. The ISME Journal 6, 554–563. doi: 10.1038/ismej.2011.135

Gaulke, A. K., Wetz, M. S., and Paerl, H. W. (2010). Picophytoplankton: A major contributor to planktonic biomass and primary production in a eutrophic, river-dominated estuary. Estuarine, Coastal and Shelf Science 90, 45–54. doi: 10.1016/j.ecss.2010.08.006

Gifford, S. M., Sharma, S., Rinta-Kanto, J. M., and Moran, M. A. (2011). Quantitative analysis of a deeply sequenced marine microbial metatranscriptome. ISME J 5, 461–472. doi: 10.1038/ismej.2010.141

Gifford, S. M., Zhao, L., Stemple, B., DeLong, K., Medeiros, P. M., Seim, H., et al. (2020). Microbial Niche Diversification in the Galápagos Archipelago and Its Response to El Niño. Frontiers in Microbiology 11, 2636. doi: 10.3389/fmicb.2020.575194

Gong, W., Hall, N., Paerl, H., and Marchetti, A. (2020). Phytoplankton composition in a eutrophic estuary: Comparison of multiple taxonomic approaches and influence of environmental factors. Environmental Microbiology 22, 4718–4731. doi: 10.1111/1462-2920.15221

Gong, W., Paerl, H., and Marchetti, A. (2018). Eukaryotic phytoplankton community spatiotemporal dynamics as identified through gene expression within a eutrophic estuary. Environmental Microbiology 20, 1095–1111. doi: 10.1111/1462-2920.14049

Grossart, H.-P., Levold, F., Allgaier, M., Simon, M., and Brinkhoff, T. (2005). Marine diatom species harbour distinct bacterial communities. Environmental Microbiology 7, 860–873. doi: 10.1111/j.1462-2920.2005.00759.x

Han, Y., Jiao, N., Zhang, Y., Zhang, F., He, C., Liang, X., et al. (2021). Opportunistic bacteria with reduced genomes are effective competitors for organic nitrogen compounds in coastal dinoflagellate blooms. Microbiome 9, 71. doi: 10.1186/s40168-021-01022-z

Haverkamp, T. H. A., Schouten, D., Doeleman, M., Wollenzien, U., Huisman, J., and Stal, L. J. (2009). Colorful microdiversity of Synechococcus strains (picocyanobacteria) isolated from the Baltic Sea. ISME J 3, 397–408. doi: 10.1038/ismej.2008.118

Hollibaugh, J. T., Gifford, S. M., Moran, M. A., Ross, M. J., Sharma, S., and Tolar, B. B. (2014). Seasonal variation in the metratranscriptomes of a Thaumarchaeota population from SE USA coastal waters. ISME J 8, 685–698. doi: 10.1038/ismej.2013.171

Isaac, A., Mohamed, A. R., and Amin, S. A. (2024). Rhodobacteraceae are key players in microbiome assembly of the diatom Asterionellopsis glacialis. Applied and Environmental Microbiology 90, e00570–24. doi: 10.1128/aem.00570-24

Jain, C., Rodriguez-R, L. M., Phillippy, A. M., Konstantinidis, K. T., and Aluru, S. (2018). High throughput ANI analysis of 90K prokaryotic genomes reveals clear species boundaries. Nature Communications 9, 5114. doi: 10.1038/s41467-018-07641-9

Johnson, Z. I., Hunt, D. E., the PICO Consortium, Wang, G., Blinebry, S., Xie, N., et al. (2025). The Piver’s Island Coastal Observatory – a decade of weekly+ observations reveal the press and pulse of a changing temperate coastal marine system. Front. Mar. Sci. 12, 1505754. doi: 10.3389/fmars.2025.1505754

Kieft, B., Li, Z., Bryson, S., Hettich, R. L., Pan, C., Mayali, X., et al. (2021). Phytoplankton exudates and lysates support distinct microbial consortia with specialized metabolic and ecophysiological traits. Proceedings of the National Academy of Sciences 118, e2101178118. doi: 10.1073/pnas.2101178118

Landa, M., Burns, A. S., Durham, B. P., Esson, K., Nowinski, B., Sharma, S., et al. (2019). Sulfur metabolites that facilitate oceanic phytoplankton–bacteria carbon flux. The ISME Journal 13, 2536–2550. doi: 10.1038/s41396-019-0455-3

Lantoine, F., and Neveux, J. (1997). Spatial and seasonal variations in abundance and spectral characteristics of phycoerythrins in the tropical northeastern Atlantic Ocean. Deep Sea Research Part I: Oceanographic Research Papers 44, 223–246. doi: 10.1016/S0967-0637(96)00094-5

Letunic, I., and Bork, P. (2024). Interactive Tree of Life (iTOL) v6: recent updates to the phylogenetic tree display and annotation tool. Nucleic Acids Research 52, W78–W82. doi: 10.1093/nar/gkae268

Li, S., Dong, Y., Sun, X., Zhao, Y., Zhao, L., Zhang, W., et al. (2024). Seasonal and spatial variations of Synechococcus in abundance, pigment types, and genetic diversity in a temperate semi-enclosed bay. Front. Microbiol. 14. doi: 10.3389/fmicb.2023.1322548

Lima-Mendez, G., Faust, K., Henry, N., Decelle, J., Colin, S., Carcillo, F., et al. (2015). Determinants of community structure in the global plankton interactome. Science 348, 1262073. doi: 10.1126/science.1262073

Lin, Y., Gifford, S., Ducklow, H., Schofield, O., and Cassar, N. (2019). Towards Quantitative Microbiome Community Profiling Using Internal Standards. Appl. Environ. Microbiol. 85, e02634–18. doi: 10.1128/AEM.02634-18

Liu, Q., Tolar, B. B., Ross, M. J., Cheek, J. B., Sweeney, C. M., Wallsgrove, N. J., et al. (2018). Light and temperature control the seasonal distribution of thaumarchaeota in the South Atlantic bight. The ISME Journal 12, 1473–1485. doi: 10.1038/s41396-018-0066-4

Luettich, R. A. J., McNinch, J. E., Paerl, H. W., Peterson, C. H., Wells, J. T., Alperin, M. J., et al. (2000). Neuse River Estuary Modeling and Monitoring Project Stage 1: Hydrography and Circulation, Water Column Nutrients and Productivity, Sedimentary Processes and Benthic-Pelagic Coupling, and Benthic Ecology. Water Resources Research Institute of the University of North Carolina. Available at: http://www.lib.ncsu.edu/resolver/1840.4/1891 (Accessed June 6, 2025).

Miller, T. R., and Belas, R. (2004). Dimethylsulfoniopropionate Metabolism by Pfiesteria-Associated Roseobacter spp. Applied and Environmental Microbiology 70, 3383–3391. doi: 10.1128/AEM.70.6.3383-3391.2004

Moran, M. A. (2015). The global ocean microbiome. Science 350, aac8455. doi: 10.1126/science.aac8455

Moran, M. A., Satinsky, B., Gifford, S. M., Luo, H., Rivers, A., Chan, L.-K., et al. (2013). Sizing up metatranscriptomics. ISME J 7, 237–243. doi: 10.1038/ismej.2012.94

Morris, R. M., Rappé, M. S., Connon, S. A., Vergin, K. L., Siebold, W. A., Carlson, C. A., et al. (2002). SAR11 clade dominates ocean surface bacterioplankton communities. Nature 420, 806–810. doi: 10.1038/nature01240

Nurk, S., Meleshko, D., Korobeynikov, A., and Pevzner, P. A. (2017). metaSPAdes: a new versatile metagenomic assembler. Genome Res. 27, 824–834. doi: 10.1101/gr.213959.116

Olm, M. R., Brown, C. T., Brooks, B., and Banfield, J. F. (2017). dRep: a tool for fast and accurate genomic comparisons that enables improved genome recovery from metagenomes through de-replication. ISME J 11, 2864–2868. doi: 10.1038/ismej.2017.126

Paerl, H. W., Crosswell, J. R., Van Dam, B., Hall, N. S., Rossignol, K. L., Osburn, C. L., et al. (2018). Two decades of tropical cyclone impacts on North Carolina’s estuarine carbon, nutrient and phytoplankton dynamics: implications for biogeochemical cycling and water quality in a stormier world. Biogeochemistry 141, 307–332.

Paerl, H. W., Rossignol, K. L., Guajardo, R., Hall, N. S., Joyner, A. R., Peierls, B. L., et al. (2009). FerryMon: Ferry-Based Monitoring and Assessment of Human and Climatically Driven Environmental Change in the Albemarle-Pamlico Sound System. Environ. Sci. Technol. 43, 7609–7613. doi: 10.1021/es900558f

Paerl, H. W., Rossignol, K. L., Hall, S. N., Peierls, B. L., and Wetz, M. S. (2010). Phytoplankton Community Indicators of Short- and Long-term Ecological Change in the Anthropogenically and Climatically Impacted Neuse River Estuary, North Carolina, USA. Estuaries and Coasts 33, 485–497. doi: 10.1007/s12237-009-9137-0

Paerl, R. W., Venezia, R. E., Sanchez, J. J., and Paerl, H. W. (2020). Picophytoplankton dynamics in a large temperate estuary and impacts of extreme storm events. Sci Rep 10, 22026. doi: 10.1038/s41598-020-79157-6

Parks, D. H., Chuvochina, M., Rinke, C., Mussig, A. J., Chaumeil, P.-A., and Hugenholtz, P. (2022). GTDB: an ongoing census of bacterial and archaeal diversity through a phylogenetically consistent, rank normalized and complete genome-based taxonomy. Nucleic Acids Research 50, D785–D794. doi: 10.1093/nar/gkab776

Parks, D. H., Imelfort, M., Skennerton, C. T., Hugenholtz, P., and Tyson, G. W. (2015). CheckM: assessing the quality of microbial genomes recovered from isolates, single cells, and metagenomes. Genome Res. 25, 1043–1055. doi: 10.1101/gr.186072.114

Peierls, B., and Paerl, H. (2010). Temperature, organic matter, and the control of bacterioplankton in the Neuse River and Pamlico Sound estuarine system. Aquat. Microb. Ecol. 60, 139–149. doi: 10.3354/ame1415

Pinckney, J. L., Paerl, H. W., Harrington, M. B., and Howe, K. E. (1998). Annual cycles of phytoplankton community-structure and bloom dynamics in the Neuse River Estuary, North Carolina. Marine Biology 131, 371–381. doi: 10.1007/s002270050330

Price, M. N., Dehal, P. S., and Arkin, A. P. (2010). FastTree 2 – Approximately Maximum-Likelihood Trees for Large Alignments. PLOS ONE 5, e9490. doi: 10.1371/journal.pone.0009490

Sanchez-Gallego, J., Curtis, N., Paerl, H. W., and Paerl, R. (2025). Shedding new light on picocyanobacteria and understudied cyanobacterial diversity in the Albemarle Pamlico Sound System, North Carolina, USA. Front. Microbiol. 16. doi: 10.3389/fmicb.2025.1539050

Sanfilippo, J. E., Nguyen, A. A., Garczarek, L., Karty, J. A., Pokhrel, S., Strnat, J. A., et al. (2019). Interplay between differentially expressed enzymes contributes to light color acclimation in marine Synechococcus. Proceedings of the National Academy of Sciences 116, 6457–6462. doi: 10.1073/pnas.1810491116

Santoro, A. E., Richter, R. A., and Dupont, C. L. (2019). Planktonic Marine Archaea. Annual Review of Marine Science 11, 131–158. doi: 10.1146/annurev-marine-121916-063141

Satinsky, B. M., Gifford, S. M., Crump, B. C., and Moran, M. A. (2013). “Chapter Twelve - Use of Internal Standards for Quantitative Metatranscriptome and Metagenome Analysis,” in Methods in Enzymology, ed. E. F. DeLong (Academic Press), 237–250. doi: 10.1016/B978-0-12-407863-5.00012-5

Scanlan, D. J., Ostrowski, M., Mazard, S., Dufresne, A., Garczarek, L., Hess, W. R., et al. (2009). Ecological Genomics of Marine Picocyanobacteria. Microbiology and Molecular Biology Reviews 73, 249–299. doi: 10.1128/mmbr.00035-08

Seemann, T. (2014). Prokka: rapid prokaryotic genome annotation. Bioinformatics 30, 2068–2069. doi: 10.1093/bioinformatics/btu153

Sharpe, G., Zhao, L., Meyer, M. G., Gong, W., Burns, S. M., Tagliabue, A., et al. (2023). Synechococcus nitrogen gene loss in iron-limited ocean regions. ISME COMMUN. 3, 1–11. doi: 10.1038/s43705-023-00314-9

Six, C., Thomas, J.-C., Garczarek, L., Ostrowski, M., Dufresne, A., Blot, N., et al. (2007). Diversity and evolution of phycobilisomes in marine Synechococcus spp.: a comparative genomics study. Genome Biol 8, 1–22. doi: 10.1186/gb-2007-8-12-r259

Stomp, M., Huisman, J., Vörös, L., Pick, F. R., Laamanen, M., Haverkamp, T., et al. (2007). Colourful coexistence of red and green picocyanobacteria in lakes and seas. Ecology Letters 10, 290–298. doi: 10.1111/j.1461-0248.2007.01026.x

Teeling, H., Fuchs, B. M., Becher, D., Klockow, C., Gardebrecht, A., Bennke, C. M., et al. (2012). Substrate-Controlled Succession of Marine Bacterioplankton Populations Induced by a Phytoplankton Bloom. Science 336, 608–611.

Urakawa, H., Martens-Habbena, W., Huguet, C., De La Torre, J. R., Ingalls, A. E., Devol, A. H., et al. (2014). Ammonia availability shapes the seasonal distribution and activity of archaeal and bacterial ammonia oxidizers in the Puget Sound Estuary. Limnology & Oceanography 59, 1321–1335. doi: 10.4319/lo.2014.59.4.1321

Vaulot, D., Eikrem, W., Viprey, M., and Moreau, H. (2008). The diversity of small eukaryotic phytoplankton (≤3 μm) in marine ecosystems. FEMS Microbiology Reviews 32, 795–820. doi: 10.1111/j.1574-6976.2008.00121.x

Vitousek, P. M., Mooney, H. A., Lubchenco, J., and Melillo, J. M. (1997). Human Domination of Earth’s Ecosystems. Science 277, 494–499. doi: 10.1126/science.277.5325.494

Wang, Z., Juarez, D. L., Pan, J.-F., Blinebry, S. K., Gronniger, J., Clark, J. S., et al. (2019). Microbial communities across nearshore to offshore coastal transects are primarily shaped by distance and temperature. Environmental Microbiology 21, 3862–3872. doi: 10.1111/1462-2920.14734

Ward, C. S., Yung, C.-M., Davis, K. M., Blinebry, S. K., Williams, T. C., Johnson, Z. I., et al. (2017). Annual community patterns are driven by seasonal switching between closely related marine bacteria. The ISME Journal 11, 1412–1422. doi: 10.1038/ismej.2017.4

Woodcroft, B. J., Aroney, S. T. N., Zhao, R., Cunningham, M., Mitchell, J. A. M., Blackall, L., et al. (2024). SingleM and Sandpiper: Robust microbial taxonomic profiles from metagenomic data. 2024.01.30.578060. doi: 10.1101/2024.01.30.578060

Zhang, J., Kobert, K., Flouri, T., and Stamatakis, A. (2014). PEAR: a fast and accurate Illumina Paired-End reAd mergeR. Bioinformatics 30, 614–620. doi: 10.1093/bioinformatics/btt593

